# Effect of Clover versus Zipping on Annular Geometry and Dynamics in Tricuspid Edge-to-Edge Repair

**DOI:** 10.64898/2026.10.25.761672

**Authors:** Magda Piekarska, Tomasz Stacel, Collin E. Haese, Dominika Sandel, Trace G. LaRue, Diego Guajardo, Austin Goodyke, Marcin Malinowski, Manikantam Gaddam, Ali Saker, Manuel K. Rausch, Tomasz A Timek

## Abstract

**Introduction:** Transcatheter edge-to-edge repair effectively reduces tricuspid regurgitation (TR), with some patients showing an additional “annuloplasty effect.”

**Methods:** Ten adult sheep underwent pulmonary artery banding to induce right ventricular failure and functional TR. Six weeks later, sonomicrometry crystals were implanted around the tricuspid annulus (TA). Two Abbott XTW clips were then placed using either a **Zipping** (n=5) or **Clover** (n=5) configuration. Echocardiographic and sonomicrometry data were collected before and after clip implantation to assess TA geometry and function.

**Results:** The model produced right ventricular dilation, dysfunction, and moderate-to-severe functional TR. In all animals, leaflet approximation with two clips significantly reduced TR severity (3.6±0.5 to 1.3±1.1; p=0.01) and septal-lateral annular diameter (29.2±2.2 to 26.7±2.9 mm; p=0.01). However, a significant reduction in septal-lateral annular diameter occurred only with the Clover technique (30±2 to 27±1 mm; p=0.001).

**Conclusions:** Both techniques reduced TR, but only the Clover configuration produced an acute annuloplasty effect by selectively decreasing septal-lateral annular diameter.

**Graphical abstract:** Does tricuspid TEER reshape the annulus? In an ovine FTR model, both Zipping and Clover techniques reduced TR, but only Clover altered septal-lateral annular geometry inducing an “annuloplasty effect.”

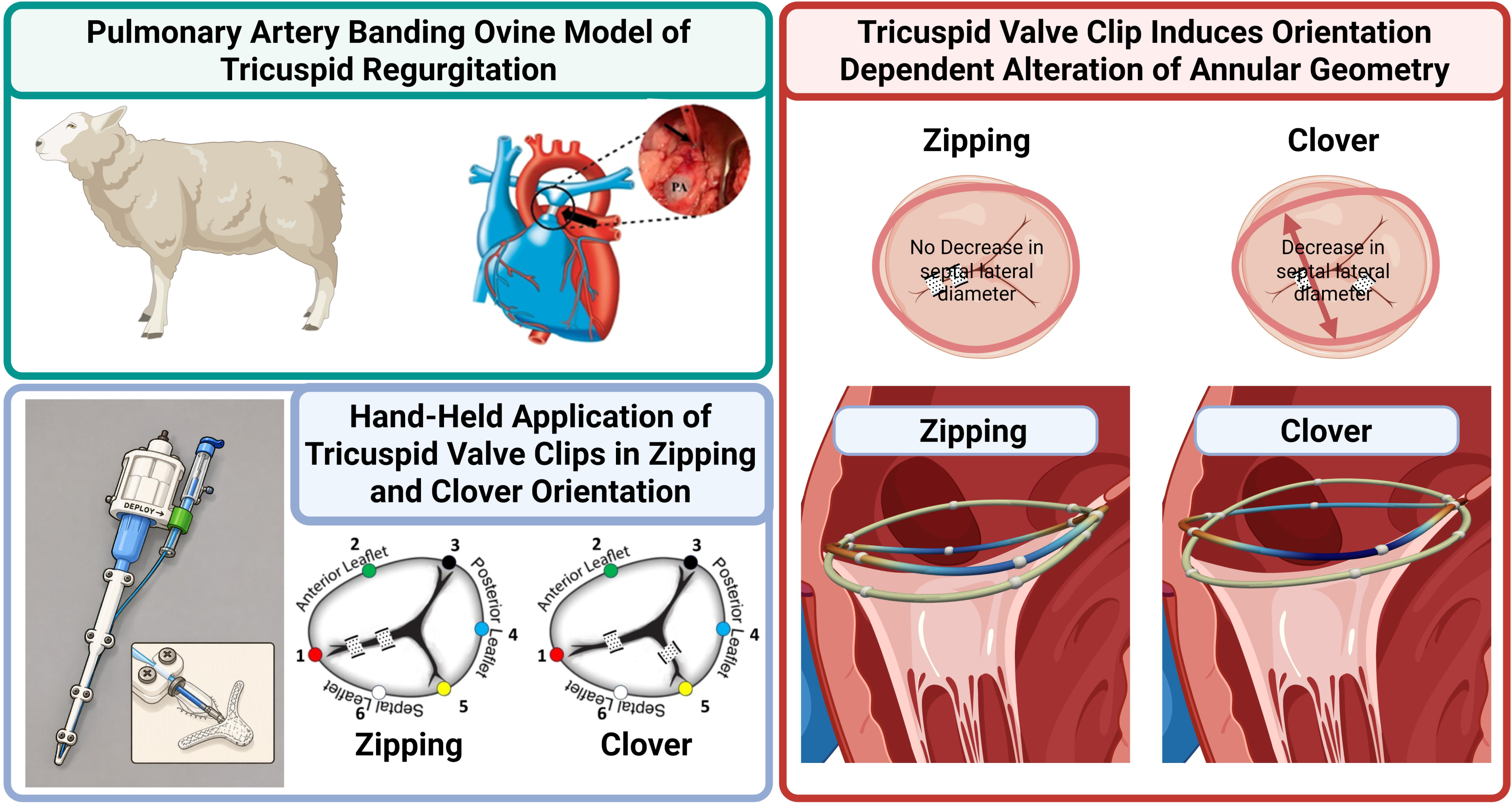

## Introduction

Severe tricuspid regurgitation (TR) is associated with worsening long-term outcomes and increasing mortality for left-sided heart disease, right ventricular (RV) insufficiency, and pulmonary hypertension.^1,2^ The most common form of TR, functional tricuspid regurgitation (FTR), is generally secondary to left sided heart disease or atrial fibrillation and associated with chamber enlargement and leaflet tethering. Dilation of the tricuspid annulus (TA) is considered a consequence of RV and right atrial remodeling and further contributes to the pathogenesis of FTR.^3,4^ Transcatheter edge-to-edge repair (TEER), or “clipping”, is an emerging technique of repair for FTR. However, treatment remains challenging due to diverse anatomy, insufficient echocardiographic imaging, and severe to torrential insufficiency.^5^ Furthermore, multiple techniques for the clipping procedure such as “Zipping” (two or more clips implanted within a single commissure) and “Clover” (clips implanted in both the antero-septal and postero-septal commissures) are offered with little data supporting superiority of either technique. In addition to decreasing TR grade, TEER may also induce annular reduction similar to surgical annuloplasty ^6–8^, yet the reports of this effect are not universal.^6^ A TEER induced “annuloplasty effect” may lead to more durable control of FTR and promote reverse remodeling, but predictors of this effect remain elusive. This knowledge gap underscores the need to re-evaluate current annuloplasty strategies and to investigate the potential annuloplasty effects of TEER.^9–12^ We set out to investigate the acute effect of Zipping and Clover TEER techniques on TR grade and tricuspid annular size, geometry, and dynamics in and ovine model of FTR.

## Methods

All sheep received humane care in compliance with the Principles of Laboratory Animal Care. The study protocols were approved by Institutional Animal Care and Use Committee Michigan State University under protocol number 202500129 approved May 14^th^, 2025. The surgical protocol for pulmonary artery banding procedure has been published previously.^10^

### Pulmonary artery banding procedure

Ten healthy Dorset adult intact male sheep had external right jugular intravenous catheter placed under local anesthesia. Animals were then anesthetized with propofol (2-5 mg/kg IV), intubated, and mechanically ventilated. General anesthesia was maintained with inhalational isoflurane (1-2.5%) and buprenorphine (0.01mg/kg iv in boluses) infused as additional maintenance anesthesia. A sterile limited left thoracotomy was made through the 4^th^ intercoastal space, and epicardial echocardiography was performed to assess biventricular function and tricuspid and mitral valve competence. The main pulmonary artery was then encircled with an umbilical tape before its bifurcation. A catheter was placed in the proximal pulmonary artery for direct pressure measurements. While monitoring systemic and pulmonary pressures, the umbilical tape was tightened down with progressive clip approximations to the brink of hemodynamic stability as described previously to at least double proximal pulmonary artery blood pressure and induce right ventricular (RV) failure and FTR.^10^ After surgical recovery, animals were monitored for 6 weeks with intermittent treatment with furosemide for evolving heart failure symptoms.

### Clipping procedure

After six weeks, the animals were returned to the operating room and anesthetized as outlined above. A median sternotomy was undertaken, and the heart was dissected from adhesions with evidence of RV dilation and dysfunction apparent (Supplementary materials - Video 1). Repeat epicardial echocardiography was performed. After full heparinization and cannulation of the right common carotid artery and jugular vein, cardiopulmonary bypass (CPB) was initiated. With the heart beating, the right atrium (RA) was opened and six 2mm sonomicrometry crystals were placed around the tricuspid annulus (Fig.1). RV and left ventricular (LV) pressure transducers were placed through the cardiac apex. Subsequently, the RA was closed, and the animal was weaned from CPB and stabilized hemodynamically. **Baseline** recordings were performed to acquire simultaneous sonomicrometry, hemodynamic, and echocardiographic data. Subsequently, CPB was reinstituted, the right atrium opened, and with the heart beating a custom designed delivery system (Fig.2) was used under direct vision to approximate the leaflets with two Abbott G4 XTW (Abbot Vascular, Menlo Park, CA) clips in two different configurations and confirmed with echocardiography (Supplementary materials - Video 2). Five sheep underwent the **Zipping (n=5)** technique, with both clips approximating the septal and anterior leaflets, and five underwent the **Clover (n=5)** technique approximating the septal leaflet to both the anterior leaflet and the posterior leaflet (Fig.3). In each configuration, the central clip approximating the septal and anterior leaflet was placed first. Following confirmation of appropriate clip placement, the right atrium was closed, and the animal weaned from CPB. After hemodynamical stabilization, sonomicrometry, hemodynamic, and echocardiographic data were re-acquired. Echocardiographic assessments were performed according to The American Society of Echocardiography guidelines and used to grade TR severity, TR max Velocity, TR Max PG, TR jet area, and vena contracta. The degree of tricuspid insufficiency was graded on a +0-4 scale as used clinically.

**Fig 1.**
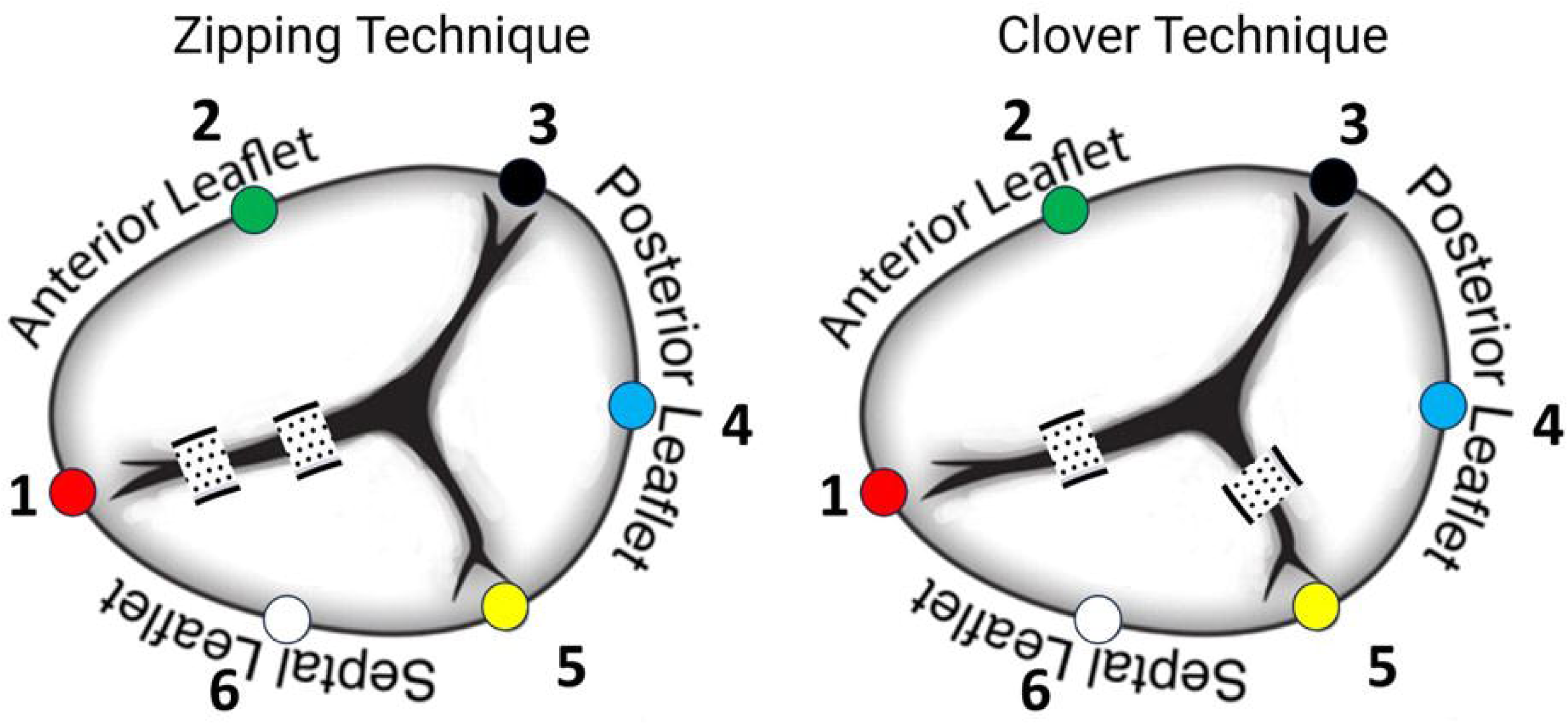
Schematic sonomicrometry crystals map with location of clips for Zipping and Clover technique. The numbers (1-6) illustrate position of implanted annular crystals: (1-2-3): anterior annulus; (3-4-5): posterior annulus; (5-6-1): septal annulus.

**Fig 2.**
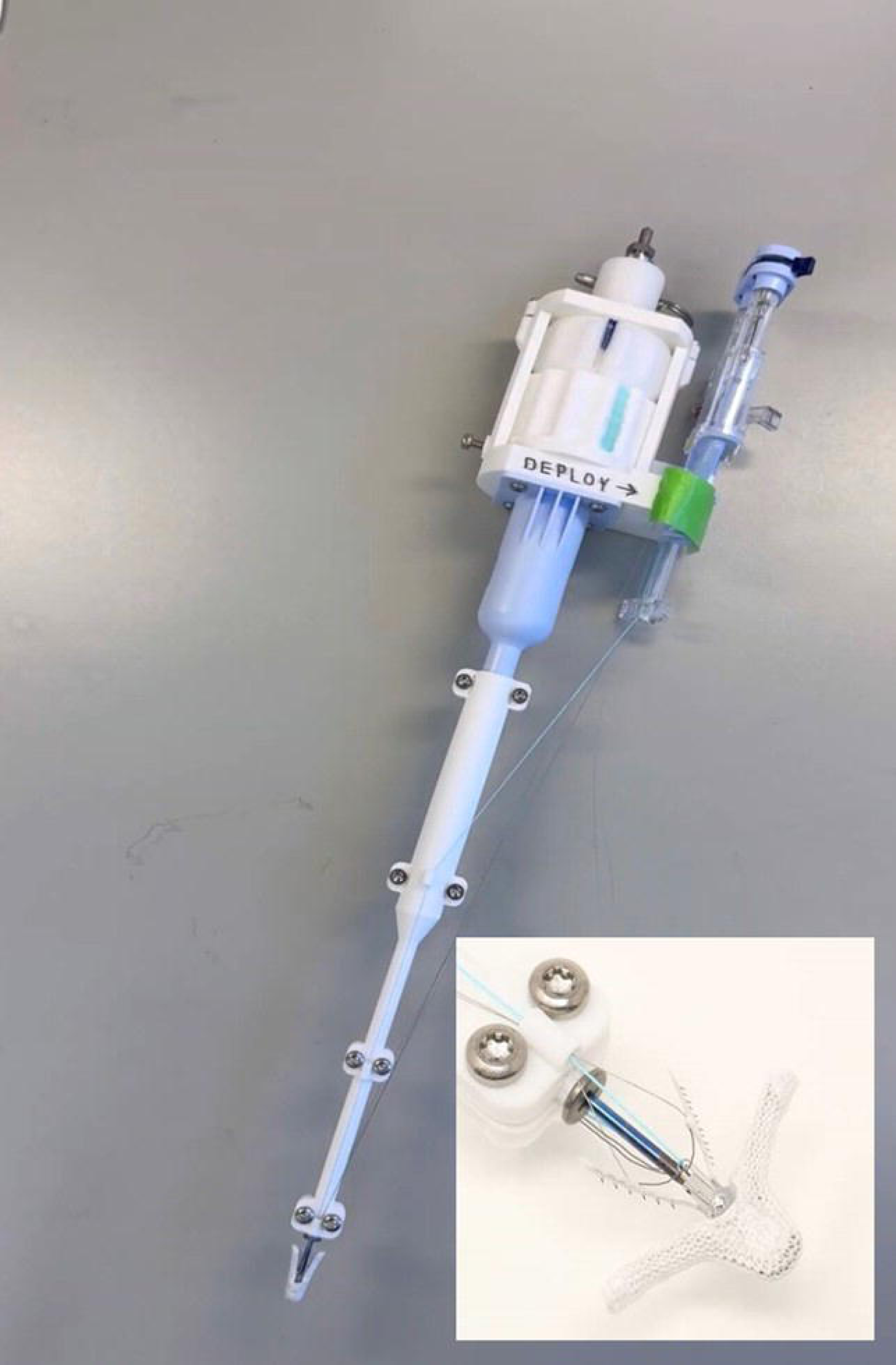
Custom designed delivery system for tricuspid impanation of an Abbott XTW TriClip through an opened atrium with direct vision. Insert illustrates the clip in an open position prior to application.

**Fig 3.**
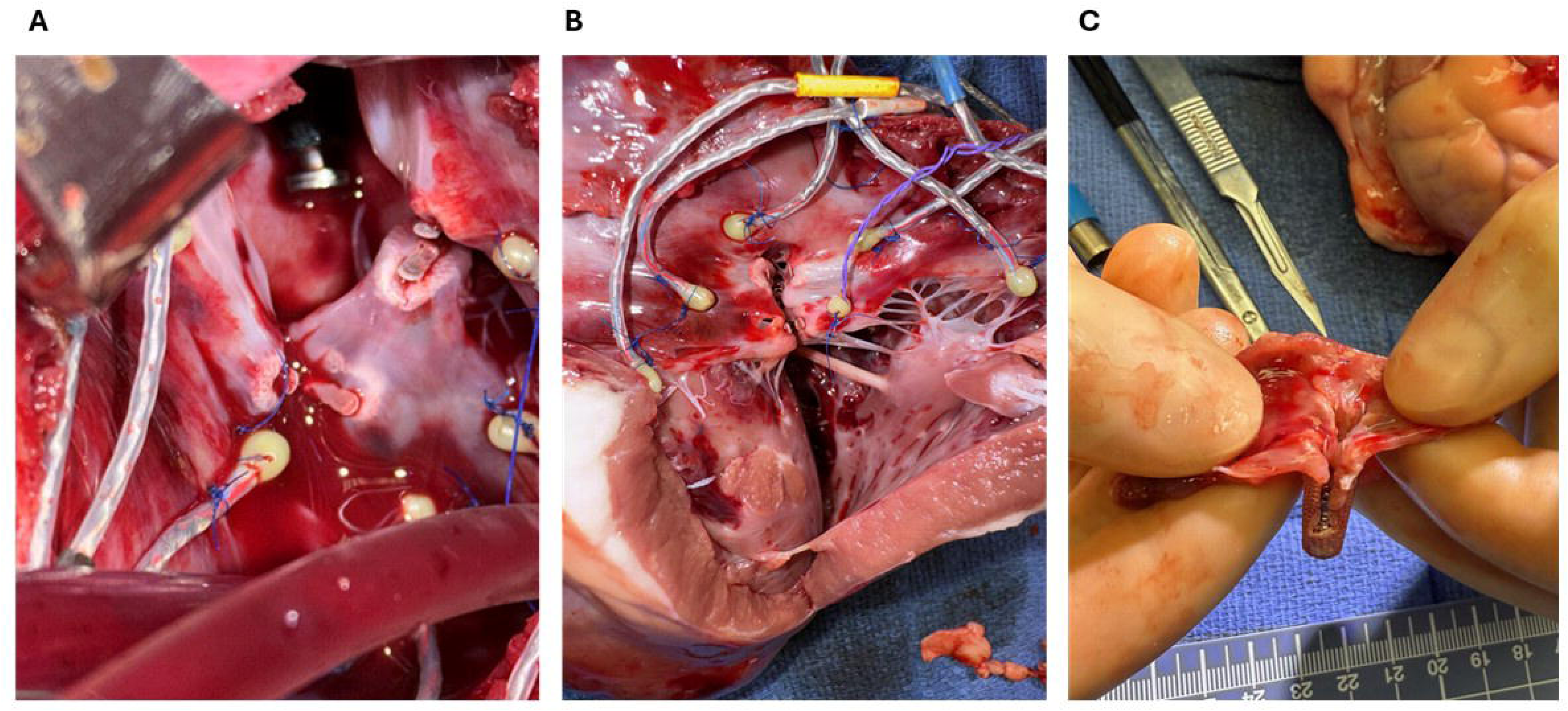
Post-mortem images of implanted sonomicrometry crystals and approximated tricuspid leaflets in a Clover (**A**) and Zipping (**B**) configuration. Panel (**C)** reveals excised leaflets confirming proper leaflet capture and full approximation.

### Data acquisition and analysis

Animals were studied with open chest and on the ventilator with respiration paused during data acquisition. Simultaneous hemodynamic, sonomicrometry, and echocardiographic data were acquired during each timepoint (Baseline, Zipping, Clover). Sonomicrometry data were acquired using a Sonomectrics Ultrasonic Measurement System DS3 (Sonometrics Inc., London, Ontario, Canada) as previously described.^13^ Sonomicrometry measurements were averaged over ten cardiac cycles in normal heart sinus rhythm before and after leaflet approximation. Data were acquired at 128 Hz with simultaneous LVP, RVP and ECG recordings. Data were analyzed with CardioSOFT Software (Sonometrics Inc.) and MATLAB code (MathWorks, Natick, Mass). Tricuspid annular area, geometry, and diameters were calculated from 3D sonomicrometry crystal coordinates as previously described.^9,10,13^ For annular strain and height analysis, the six sonomicrometry crystals sutured around the tricuspid annulus provided three-dimensional marker coordinates at each experimental condition. For each heart, marker coordinates were fit with a smooth spline to create a continuous mathematical representation of the annulus, as previously described by our group.^14,15^ This was done separately at end-diastole and end-systole and separately for the Clipping and Zipping groups. Baseline annular geometry, obtained before TEER, served as the reference configuration, matched by cardiac phase (baseline end-diastole compared with post-intervention end-diastole, and likewise for end-systole). Annular strain was calculated along the length of the annulus as the local percent change in distance between corresponding points on the deformed (Clipping or Zipping) annulus relative to baseline, with positive strain indicating lengthening and negative strain indicating shortening of the annular tissue. Relative annular height was calculated as the change in distance of each point along the annulus from the annular plane, comparing the Clipping or Zipping configuration with the baseline at the same cardiac phase. Strain and height were displayed as color-coded maps along the annular circumference for the Clipping and Zipping groups separately at end-diastole and end-systole. Negative or compressive strains imply that tissue was contracted whereas positive or tensile strains imply that tissue is stretched.^14,15^

### Statistical analysis

Data are presented as mean ± standard deviation. Measurements were compared using a paired Student two-tailed t-test, or for data not meeting normality assumptions, a non-parametric equivalent Wilcoxon signed-ranked test was used. A p value <0.05 was considered significant. All statistical analyses were performed using MedCalc Statistical Software, version 20.114.

## Results

All animals survived the experimental protocol. Table 1 summarizes animal weight, TR, and pulmonary artery pressures before and after pulmonary banding and at six-week follow-up. Animal weights in Zipping and Clover groups were alike before pulmonary banding and after six weeks. Both cohorts underwent similar hemodynamic RV insult as evidenced by comparable systolic pulmonary artery pressure immediately after banding and at 6-week follow-up. Zipping and Clover animals developed similar level of moderately severe FTR (3.3±0.4 vs 3.8±0.4, p=0.12). Relative to pre-clipping baseline, leaflet approximation in all animals reduced TR grade from 3.6±0.5 to 1.3±1.1 (p=0.01) while reducing the echocardiographic RV1 diameter from 26.6±2.8 to 20.7±4.8 mm (p=0.01). All animals achieved TR reduction of moderate or less except one animal in the Zipping group which had continued severe TR. Leaflet approximation achieved comparable reduction in TR in Clover (3.8±0.4 to 1.1±0.5; p=0.002) and Zipping (3.3±0.4 to 1.5±1.5 p=0.06) but this reached statistical significance only in the Clover animals.

**Table 1.**
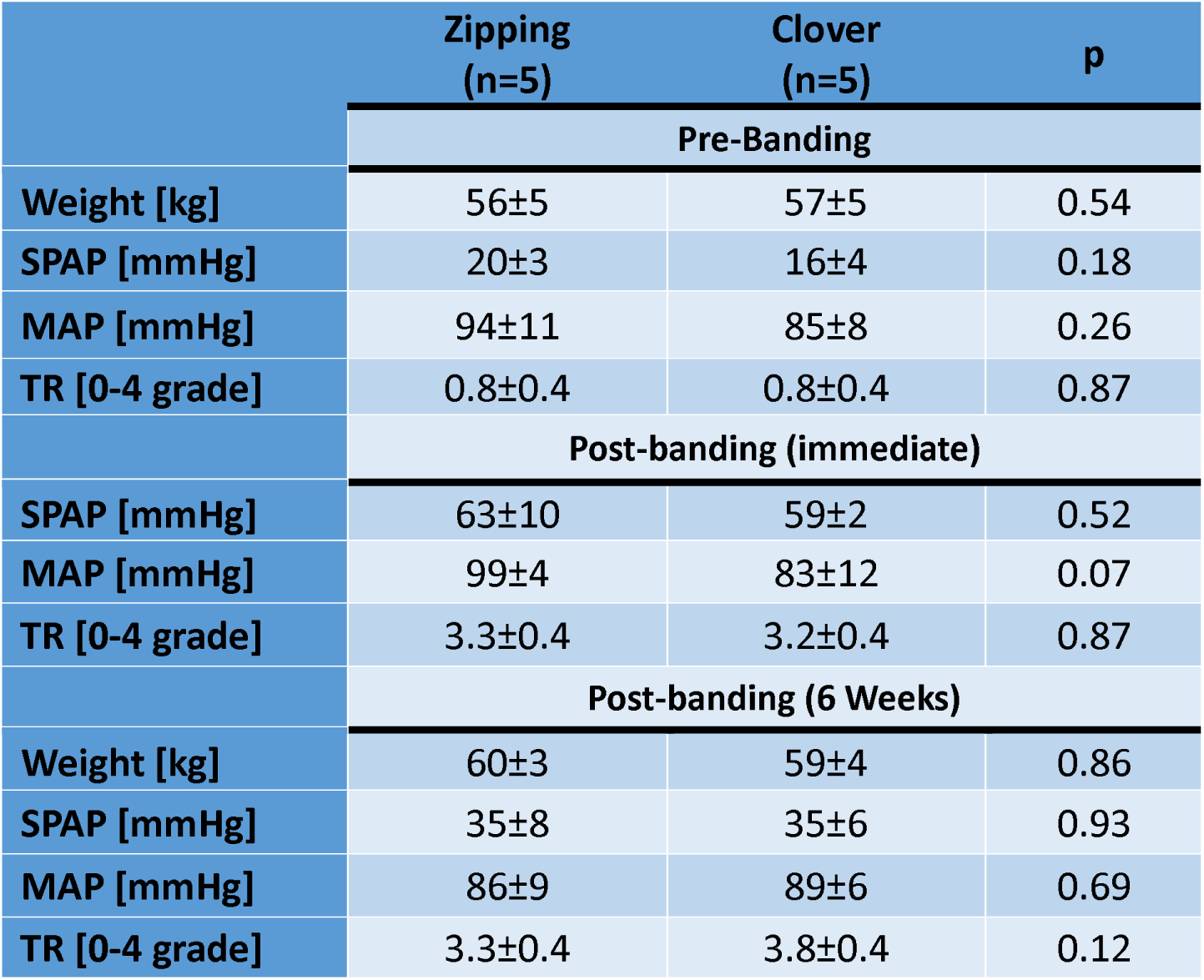
Hemodynamic data. SPAP= systolic pulmonary artery pressure, MAP= mean systemic arterial pressure, TR= tricuspid regurgitation. Values are reported as mean ± one standard deviation; * p < 0.05 by Student’s t-test for dependent observations.

|  | **Zippling**<br>**(n=5)** | **Clover**<br>**(n=5)** | **p** |
| --- | --- | --- | --- |
|  | **Pre-Banding** |  |  |
| **Weight [kg]** | 56±5 | 57±5 | 0.54 |
| **SPAP [mmHg]** | 20±3 | 16±4 | 0.18 |
| **MAP [mmHg]** | 94±11 | 85±8 | 0.26 |
| **TR [0-4 grade]** | 0.8±0.4 | 0.8±0.4 | 0.87 |
|  | **Post-banding (immediate)** |  |  |
| **SPAP [mmHg]** | 63±10 | 59±2 | 0.52 |
| **MAP [mmHg]** | 99±4 | 83±12 | 0.07 |
| **TR [0-4 grade]** | 3.3±0.4 | 3.2±0.4 | 0.87 |
|  | **Post-banding (6 Weeks)** |  |  |
| **Weight [kg]** | 60±3 | 59±4 | 0.86 |
| **SPAP [mmHg]** | 35±8 | 35±6 | 0.93 |
| **MAP [mmHg]** | 86±9 | 89±6 | 0.69 |
| **TR [0-4 grade]** | 3.3±0.4 | 3.8±0.4 | 0.12 |

Based on sonomictrometry determined 3D crystal coordinates, annular septal-lateral diameter of all study animals decreased from 29.2±2.2 mm to 26.7±2.9 mm (p=0.01) after leaflet approximation representing an approximate 8% reduction in annular size. Neither tricuspid annular area (857±132 versus 851±157 mm^2^, p=0.9) nor perimeter (107±10 vs 110±13 mm, p=0.48) decreased with clipping. Sonomicrometry based calculations of tricuspid annular geometry and size at end-diastole and global and regional annular dynamics with Zipping and Clover repairs are provided in Table 2. Due to dislocation of one of the clips in the Zipping group, only 4 animals in this group were available for analysis. TA area or perimeter did not change from Baseline for either Zipping or Clover. Furthermore, annular area did not change throughout the cardiac cycle with either of the two techniques as demonstrated in Fig.4. However, TA septal-lateral diameter was significantly reduced with Clover (30±1.5 to 27±0.8 mm, p=0.001) but this was not observed with Zipping (28±2.5 to 26±4.6 mm, p=0.36). Clover repair was associated with significant reduction in length of the posterior annular segment and increased size of the antero-septal to postero-septal commissure distance. Zipping significantly reduced the dynamic contraction of the entire annular perimeter and the contraction of the anterior and posterior segments of the annulus.

**Fig 4.**
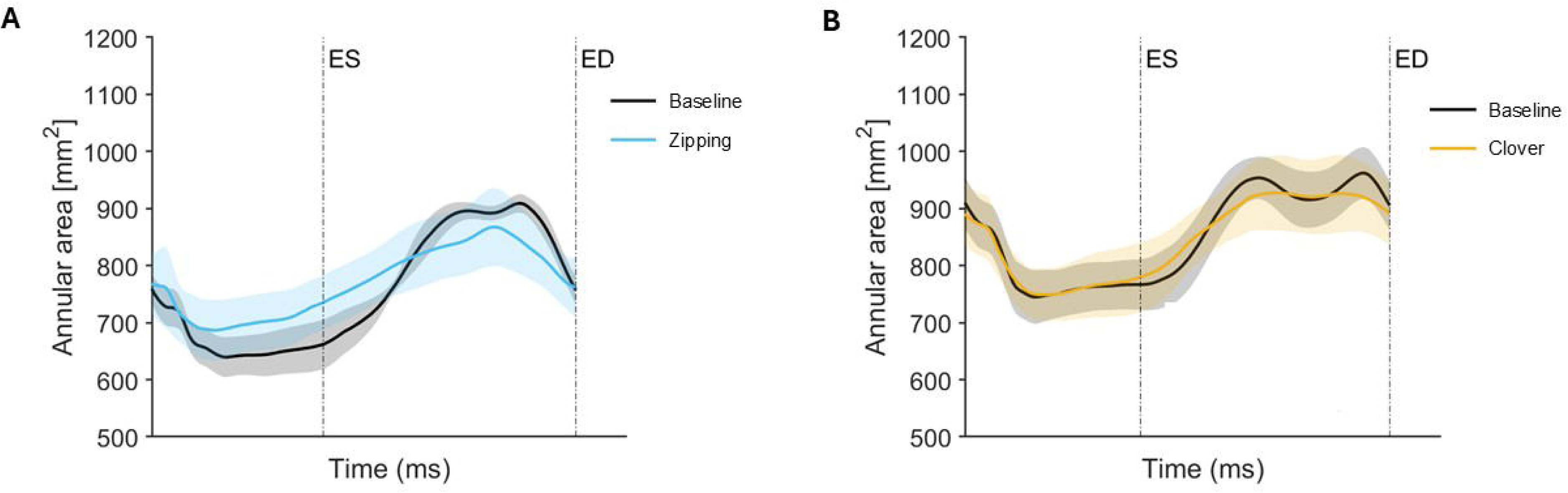
Group mean tricuspid annular area for Zipping (blue, panel **A**) and Clover (orange, panel **B**) versus respective Baseline (black) throughout the cardiac cycle. ES= end-systole, ED= end-diastole. Shaded areas represent ± one standard deviation.

**Table 2.** Sonomicrometry data. TAA= tricuspid annular area, TAP= Tricuspid annular perimeter. Values are reported as mean ± one standard deviation; *p<0.05 by Student’s t-test for dependent observations.

|  |  | Baseline | Zippering (n=4) | p | Baseline | Clover (n=5) | p |
| --- | --- | --- | --- | --- | --- | --- | --- |
| TAA (mm<sup>2</sup>) |  | 768±84 | 779±158 | 0.89 | 928±123 | 909±146 | 0.78 |
| TAP (mm) |  | 102±9 | 108±18 | 0.47 | 111±9 | 111±9 | 0.98 |
| TAA Contraction (%) |  | 31±12 | 22±13 | 0.14 | 25±9 | 22±9 | 0.39 |
| TAP Contraction (%) |  | 14±6 | 9±6 | 0.04* | 13±4 | 10±4 | 0.31 |
| TA Diameter (mm) | AP (1-4) | 32±3 | 32±4 | 0.79 | 32±4 | 34±5 | 0.43 |
|  | SL (2-6) | 28±3 | 26±5 | 0.36 | 30±2 | 27±1 | 0.001* |
| Eccentricity | AP/SL | 0.65±0.2 | 0.67±0.1 | 0.81 | 0.76±0.2 | 0.55±0.1 | 0.16 |
| Regional TAP (mm) | Anterior (1-2-3) | 33±13 | 34±9 | 0.81 | 39±6 | 39±7 | 0.71 |
|  | Posterior (3-4-5) | 30±5 | 32±10 | 0.41 | 33±9 | 30±9 | 0.02* |
|  | Septal (5-6-1) | 39±9 | 42±13 | 0.31 | 40±12 | 42±11 | 0.08 |
| Intercomissure distance (mm) | AS-AP (1-3) | 26±7 | 25±7 | 0.38 | 29±4 | 32±4 | 0.17 |
|  | AP-PS (3-5) | 26±6 | 26±10 | 0.91 | 28±6 | 26±7 | 0.15 |
|  | AS-PS (5-1) | 32±4 | 33±5 | 0.35 | 32±7 | 35±6 | 0.04* |
| Regional TAP Contraction (%) | Anterior (1-2-3) | 18±6 | 12±6 | 0.02* | 15±5 | 14±4 | 0.67 |
|  | Posterior (3-4-5) | 16±8 | 10±7 | 0.03* | 16±5 | 12±4 | 0.32 |
|  | Septal (5-6-1) | 14±9 | 11±9 | 0.32 | 14±5 | 11±5 | 0.20 |

Annular 3D geometry represented by annular height along the entire length of the perimeter at end-diastole is summarized for Zipping and Clover relative to their respective Baseline in Fig.5. These reconstructions reveal elevation of segments of the septal and lateral annulus with leaflet approximation with deeper nadirs near the commissures accentuating the “saddle” shape of the tricuspid annulus. Annular strain representing regional contractility for both approximations at end diastole is depicted in Fig.6. Annular strain was accentuated in sepal and lateral regions of the annulus relative to Baseline in both approximations but appeared to be more pronounced with the Clover technique.

**Fig 5.**
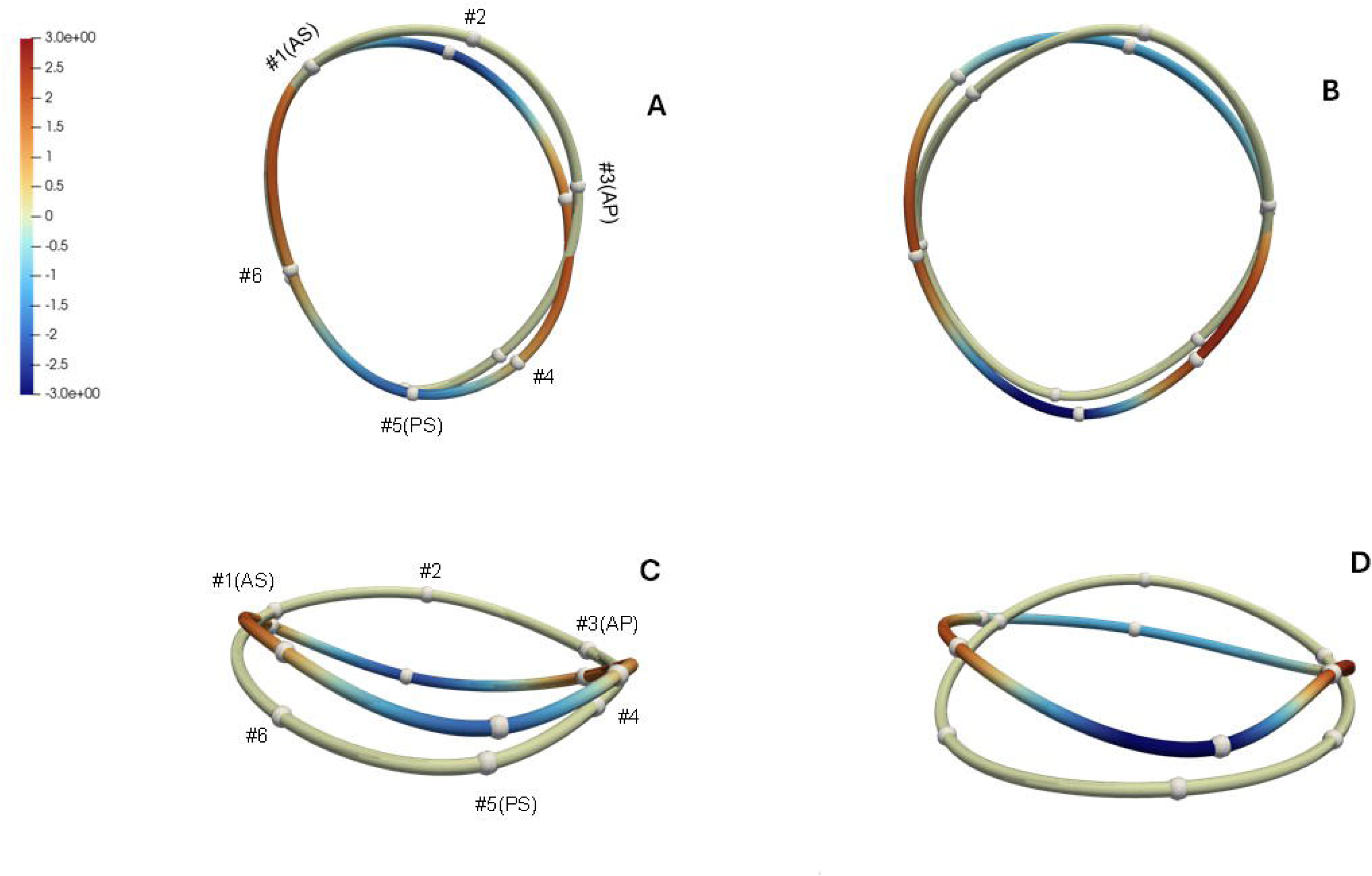
Group mean annular height at end-diastole at baseline (reference) and with clipping (colored figure) for Zipping (**A, C**) and Clover (**B, D**). Panels **A** and **B** are 2D projections from the atrial view, while panels **C** and **D** represent corresponding posterior views. Scale represents regional increase (red) or reduction (blue) in annular height from -3 to 3 mm.

**Fig 6.**
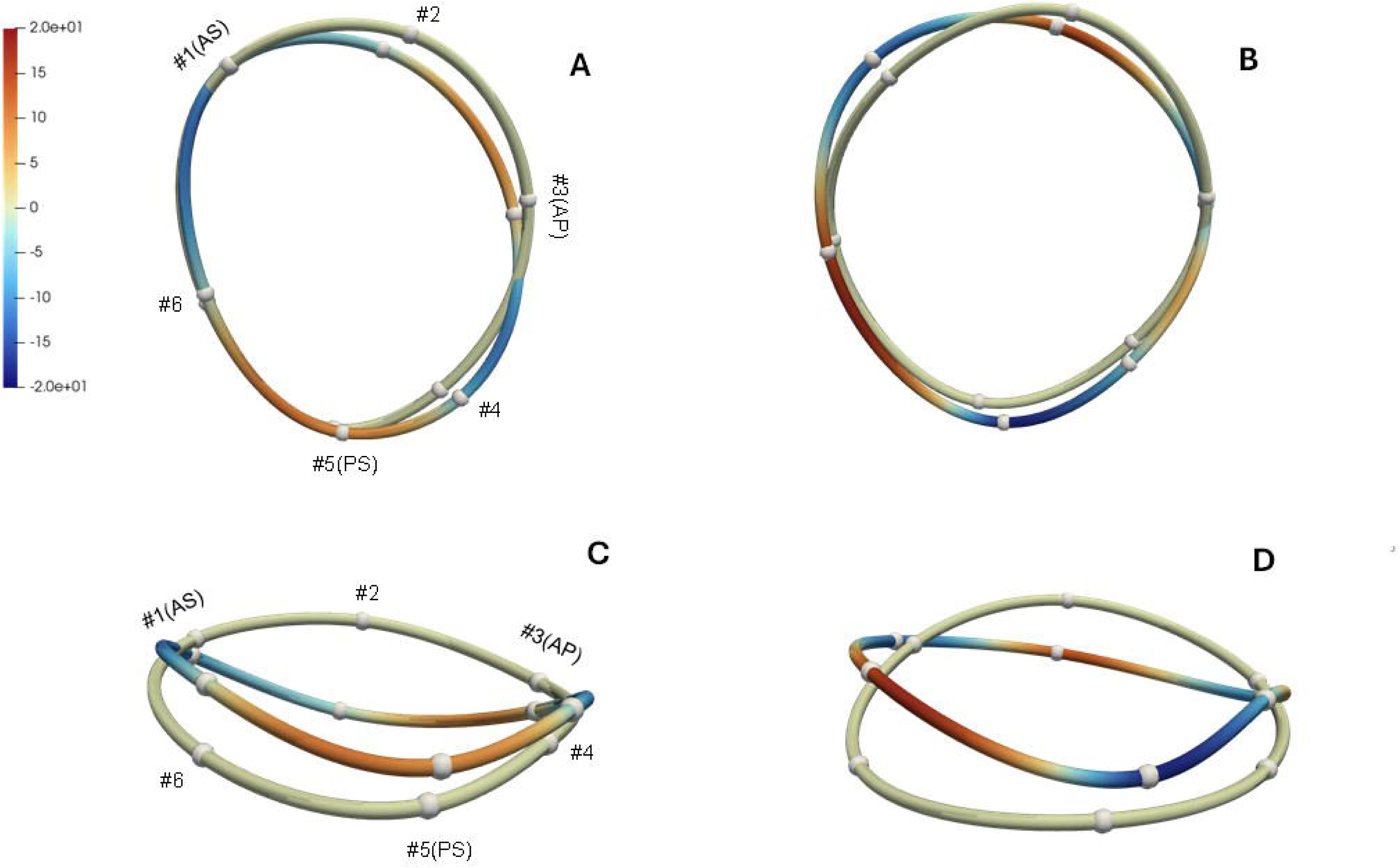
Group mean annular strain along the tricuspid annular circumference at end-diastole at baseline (reference) and with clipping (colored figure) for Zipping (**A, C**) and Clover (**B, D**). Panels **A** and **B** are 2D projections from the atrial view, while panels **C** and **D** represent corresponding posterior views. Scale represents regional contraction (red) or lengthening (blue) of the annulus from 20 to -20%.

## Discussion

Our ovine model of pulmonary banding resulted in moderately severe functional TR with right ventricular failure mimicking the clinical disease. Leaflet approximation through the direct application of a clip significantly reduced TR grade and septal-lateral annular diameter without affecting annular area or perimeter. In both Zipping and Clover leaflet approximation techniques, we observed a comparable decline in tricuspid regurgitation, but only the Clover approximation achieved significant annular reduction in the septal-lateral dimension. These data suggest a differential influence of approximating clip configuration on the “annuloplasty effect” of TEER.

Both techniques of leaflet approximation utilized in our study achieved a therapeutic effect of reducing FTR with 8/9 (88%) of animals achieving moderate or less insufficiency after clipping, consistent with reported clinical data.^16^ We used direct vision and an open right atrium to place clips in a consistent location in each group assuring full leaflet capture and no entanglement of the subvalvular apparatus. However, we still experienced a technical failure with one clip dislodging when attempting the Zipping technique. In TRILUMINATE PIVOTAL study^17^ “device success” with TEER was 88.9% with two-year rate of single-leaflet device detachment at 5.6%^18^ and 84% of patients having moderate or less TR. Similar results were reported in Transatlantic TRILUMINATE Trial^19^ with reduction of TR of at least one grade achieved in 85.4% of subjects and maintained in 75% of the study population. Echarte-Morales group^20^ compared zipping and clover techniques and reported greater residual TR with the clover technique, but these patients also had more complex and challenging TV anatomy. Clinical outcomes, however, were similar between both techniques. In the current study, one animal in the Zipping group had persistent severe TR and as such, significant TR reduction was only observed in the Clover group. However, the grade of TR after clipping was comparable between the two groups and the lower number of animals in the Zipping group may have contributed to this lack in significance.

The effect of leaflet approximation on annular geometry was first demonstrated in mitral TEER more than a decade ago.^21,22^ The potential annular remodeling effect of tricuspid leaflet approximation was first suggested by Hausleiter who postulated that clover type approximation would be most effective.^23^ *Ex vivo* porcine tricuspid studies further suggested that mid-leaflet grasping rather than commissural approximation may be more advantageous.^24^ The clinical effect of tricuspid annular size reduction, the “annuloplasty effect”, through T-TEER was first demonstrated by Russo and colleagues who found that 62% of patients undergoing tricuspid leaflet approximation experienced annular diameter reduction.^8^ Mean annular diameter reduction was approximately 5.2 mm (11%) and these results were corroborated by findings of a smaller Italian study.^25^ The acute “annuloplasty effect” was reported by Cammalleri and colleagues who performed 3D transesophageal echocardiography immediately prior and after T-TEER with the TriClip.^7^ These investigators found a significant median septal-lateral annular reduction of approximately 5 mm after leaflet approximation which was independent of procedural success. Our study demonstrated a significant acute reduction in SL diameter of approximately 8% in all clipped animals, which is consistent with clinical reports. However, we also found a differential effect of Clover versus Zipping on annular size reduction with only the former achieving a significant effect. These data are supported by Von Stein et al. who in a retrospective clinical study based on 3D transesophageal echocardiography found superior annular reduction with clover versus zipping strategy yet the effect on TR grade was similar.^6^ Furthermore, our findings are in line with T-TEER annular effects predicted by Hausleiter.^23^ There is a paucity of comparative data between T-TEER approximating strategies in literature and clinical studies have employed a myriad of clipping locations and number of deployed clips making direct comparisons difficult.^8,20,26,27^ Computational modeling studies have suggested that central leaflet edge clipping with larger clips may accentuate annular reduction which is mainly driven by decrease in septal-lateral diameter.^28^ Furthermore, the greatest increase in cardiac output in an *ex vivo* porcine model was observed with a Clover like configuration.^24^ Recent one-year results from the PASTE registry of over 1000 T-TEER patients suggest that clover approximation was associated with improved echocardiographic outcomes. Procedural annuloplasty-oriented T-TEER report by Shuvy^29^ suggested that the approximating strategy should be tailored to leaflet anatomy, yet Clover approximation would be expected to achieve greater annuloplasty effect in each scenario. Zipping approximation, in our hands, also resulted in significant reductions in tricuspid annular perimeter contraction, further indicating a more beneficial effect of Clover approximation. However, clinical confirmation of our experimental work on the differential “annuloplasty effect” of Clover versus Zipping remains to be determined.

In our study we did not observe a change in annular area or perimeter with leaflet approximation although this has been reported in clinical studies using both transthoracic^8,25^ and transesophageal echocardiography.^6–8,25^ This discrepancy may be related to anatomic considerations and imaging techniques as the complex three-dimensional shape of the tricuspid annulus^4^ does not lend itself easily to precise assessment. Sonomicrometry crystals permit tracking of discreet points on the tricuspid annulus throughout the cardiac cycle with sub-millimeter spatial resolution^9^ which may be more difficult with echocardiography. Although area change was not apparent, our data illustrated significant annular deformation with both Clover and Zipping techniques. The three-dimensional shape of the tricuspid annulus became accentuated with leaflet approximation which may in turn have a positive effect of stress distribution and flow dynamics. This was seen in the elevation of annular segments in the areas of approximation suggesting annular folding with T-TEER. The girdling effect of leaflet clipping was also demonstrated in our annular strain data revealing regional differences in annular length at end-diastole induced by T-TEER. The annular location specific changes elucidated by these data may better guide direction of leaflet approximation to maximize the “annular effect” of this intervention. These results also emphasize the need to evaluate the influence of T-TEER on the tricuspid annulus in the context of its complex three-dimensional shape and dynamic physiology rather than size alone. Our data suggest that tricuspid annular size, geometry, and dynamics are all influenced by leaflet approximation.

## Conclusions

Percutaneous tricuspid leaflet edge-to-edge approximation holds the promise of improved therapy for a difficult group of patients with advanced FTR, yet little is known regarding the effect of T-TEER on annular physiology. Our experimental study in sheep with chronic FTR found that leaflet approximation with two clips significantly reduced FTR and annular septal-lateral diameter. Clover approximation was more effective in reducing annular size while minimizing reductions in contraction in comparison to Zipping approximation. These data may better guide clinical tailoring of TEER to maximize the “annuloplasty effect” of the procedure.

### Limitations

Extrapolation of our experimental data to the clinical environment should be performed with caution due to several important limitations of our study. Our ovine model of FTR and RV dysfunction was induced by pulmonary banding over six weeks which departs from clinical evolution of FTR over years. Chamber dilation and level of torrential TR often seen in clinical studies^7,25,30^ was not matched by our model, and therefore the effects of our intervention may be overestimated. Furthermore, we implanted the clips through and opened atrium while on cardiopulmonary bypass adding a physiologic insult normally not seen clinically. However, the beating heart technique minimized any untoward effect on RV function and the opened atrium permitted precise and reproducible clip placement that may be more difficult with image guided approximations. Leaflet anatomy was very consistent in our animals representing uniformly Type 1 anatomy, but up to 30% of patients may have Type IIIb anatomy which may contribute to different effects of T-TEER than observed in our study. Crystal sonomicrometry offers a precise measurement platform but requires suturing of crystals to the tricuspid annulus which may in turn affect annular physiology. However, this effect is minimal with no differences in mitral annular physiology reported between implanted miniature wireless tantalum markers^31^ and sonomicrometry crystals. The low number of animals in the study presents yet another limitation especially in the Zipping group, although the paired design of the study provided additional statistical power. Our results offer only data on the acute intra-procedural effect of leaflet approximation and any reference to distant outcomes should be avoided. We used only castrated males in the study and therefore the effect of sex, an important physiologic determinant, on the study end-points cannot be assessed.

## Clinical relevance

Transcatheter edge-to-edge repair has been shown to be a successful strategy in reducing TR grade in patients with function tricuspid regurgitation, with evidence of an “annuloplasty effect” demonstrated in some patients. In comparing Zipping and Clover techniques of leaflet approximation, in this study only Clover technique achieved a significant “annuloplasty effect”. These data may better guide clinical application of T-TEER to optimize annular remodeling and long-term control of FTR.

## Acknowledgements

Magda Piekarska and Tomasz Stacel are Advanced Translational Research Fellows in Cardiothoracic Surgery at Corewell Health.

## Disclosures

Manuel K Rausch has a speaking arrangement with Edwards Lifesciences. The authors have no other disclosures associated with this manuscript.

**No human studies were carried out by the authors for this article.**

**All sheep in this study received humane care in compliance with the Principles of Laboratory Animal Care. The study protocols were approved by Institutional Animal Care and Use Committee Michigan State University under protocol number 202500129 approved May 14^th^, 2025.**

## Abbreviations

AP: antero-posterior
CPB: cardiopulmonary bypass
CVP: central venous pressure
ECG: electrocardiogram
ED: end-diastole
ES: end-systole
FTR: functional tricuspid regurgitation
LV: left ventricle
PAB: pulmonary artery banding
PS: postero-septal
RA: right atrium
RV: right ventricle
SL: septal-lateral
TA: tricuspid annulus
TAA: tricuspid annulus area
TAP: tricuspid annulus perimeter
TEER: transcatheter edge-to-edge repair
TR: tricuspid regurgitation
TS: tricuspid stenosis

**Supplementary Materials - Video 1.** Intraoperative video of the beating heart after six weeks of pulmonary banding. Visible is remarkable right ventricular enlargement and dysfunction induced by chronic pressure overload and functional tricuspid regurgitation.

**Supplementary Materials - Video 2.** Intra-operative video illustrating implanted sonomicrometry crystals and approximated tricuspid leaflet on the beating heart while on cardiopulmonary bypass.

